# STEAM: Spatial Transcriptomics Evaluation Algorithm and Metric for clustering performance

**DOI:** 10.1101/2025.02.17.636505

**Authors:** Samantha Reynoso, Courtney Schiebout, Revanth Krishna, Fan Zhang

## Abstract

**Motivation:** Spatial transcriptomic technologies allow researchers to explore the diversity and specificity of gene expression within their original tissue structure. Accurately identifying regions that are spatially coherent in both gene expression and physical tissue structures is an emerging topic, but challenging due to the lack of ground truth labels which renders complicating validation of clustering consistency and reproducibility. This highlights a need for a computational evaluation framework to rigorously and unbiasedly assess clustering performance.

**Results:** To address this gap, we propose STEAM (Spatial Transcriptomics Evaluation Algorithm and Metric), a user-friendly computational pipeline designed to evaluate the consistency and reliability of clustering results by leveraging machine learning classification and prediction methods, with the goal of maintaining the spatial proximity and gene expression patterns within clusters. We benchmarked STEAM on various public datasets, spanning multi-cell to single-cell resolution, as well as spatial transcriptomics and proteomics. The results highlighted its robustness and generalizability through comprehensive statistical evaluation metrics, such as Kappa score, F1 score, accuracy, and adjusted rand index. Notably, STEAM supports multi-sample training, enabling cross-replicate clustering consistency assessment. Moreover, STEAM provides practical guidance by comparing clustering results across multiple approaches; here, we evaluated four different methods, including spatial-aware and spatial-ignorant approaches. In summary, we believe that STEAM provides researchers a promising tool for evaluating clustering robustness and benchmarking clustering performance for spatial omics data, offering valuable insights to drive reproducible discoveries in spatial biology.

**Availability and implementation:** Source code and the R software tool STEAM are available from https://github.com/fanzhanglab/STEAM.

## Introduction

The recent advancement in spatial transcriptomics and other spatial omics technologies have made it possible to investigate cellular heterogeneity and the specificity of gene expression while preserving their spatial organization within tissues (Larsson et al., 2021; Moses and Pachter, 2022). The development of computational methods plays a critical role in analyzing spatial omics data, with unsupervised clustering methods being widely applied to uncover biologically meaningful groups for further interpretation (Miller et al., 2021). Several computational clustering methods have been developed, including spatial-ignorant methods like gene expression-driven Louvain (Blondel et al., 2008) and Leiden clustering (Traag et al., 2019), and spatial-aware methods such as BayesSpace (Zhao et al., 2021), SpaceFlow (Ren et al., 2022), DR.SC (Liu et al., 2022), and STew (Guo et al., 2024). The spatial-aware clustering methods aim to address the challenge of identifying clusters that preserve both gene expression patterns and spatial coherence. However, the methodological differences and lack of unbiased evaluation metrics pose a challenge in evaluating the robustness of identified cell type clusters. This challenge is further exacerbated by the frequent absence of ground truth cell type labels in real datasets, particularly in poorly characterized or diseased tissues, making rigorous validation even more difficult. Given the importance of cluster consistency in spatial omics analysis, there is a need for a rigorous, quantitative, and unified computational framework to systematically evaluate the consistency and reliability of clustering results.

While standard statistical metrics, such as Silhouette score, LISI (Local Inverse Simpson’s Index) (Korsunsky et al., 2019), and PAS (percentage of abnormal spots) score (Shang and Zhou, 2022) are commonly used to evaluate clustering quality for standard single-cell transcriptomics, they fall short in capturing the classification capability and predictability of clusters in the spatial context, especially when considering spatial proximity and gene expression variation in spatial omics. Moreover, assessing cluster reproducibility across multiple replicates generated from spatial omics presents a greater challenge. Unlike standard single-cell transcriptomics, where clusters are often well-defined based on transcriptomic profiles alone, spatial omics introduce additional complexity due to spatially correlated gene expression patterns and variations in tissue microenvironments. Thus, developing a rigorous computational framework tailored for spatial omics will empower researchers to more accurately assess the strengths and limitations of clustering results, leading to more reliable, reproducible, and biologically meaningful clusters in downstream analyses.

To address this, we propose STEAM, a Spatial Transcriptomics Evaluation Algorithm and Metric for clustering performance, an user-friendly strategy designed to evaluate the consistency of clustering algorithms and the reliability of cell annotations in spatial omics data. Our hypothesis is that if clusters are robust and consistent across tissue regions, selecting a subset of cells or spots within a cluster should enable accurate prediction of cell type annotations for the remaining cells within that cluster, due to spatial proximity and gene expression covarying patterns. STEAM integrates various machine learning classification models to assess the prediction accuracy and consistency of clusters, combined with unbiased statistical evaluation metrics. We demonstrated the generalizability and scalability of STEAM across various spatial omics platforms, spanning multi-cell to single-cell resolution, as well as spatial transcriptomics and proteomics. Furthermore, we used STEAM to evaluate the performance of spatial-aware and spatial-ignorant clustering methods, offering researchers a valuable tool for more informed result interpretation to guide toward biologically robust results.

## Method

### Overview of STEAM

An overview of STEAM workflow is illustrated in **Figure 1**. STEAM takes in a spatial transcriptomic dataset, either single-sample or multi-sample, along with spatial coordinates and labels provided by the clustering or annotation. STEAM then evaluates the method’s reliability by measuring prediction consistency across data splits using machine learning models, for example, Random Forest (RF) and Support Vector Machine (SVM). Investigators can use this output to evaluate the consistency of cell type annotation results, compare methods, and make informed decisions for downstream analysis.

**Figure 1.**
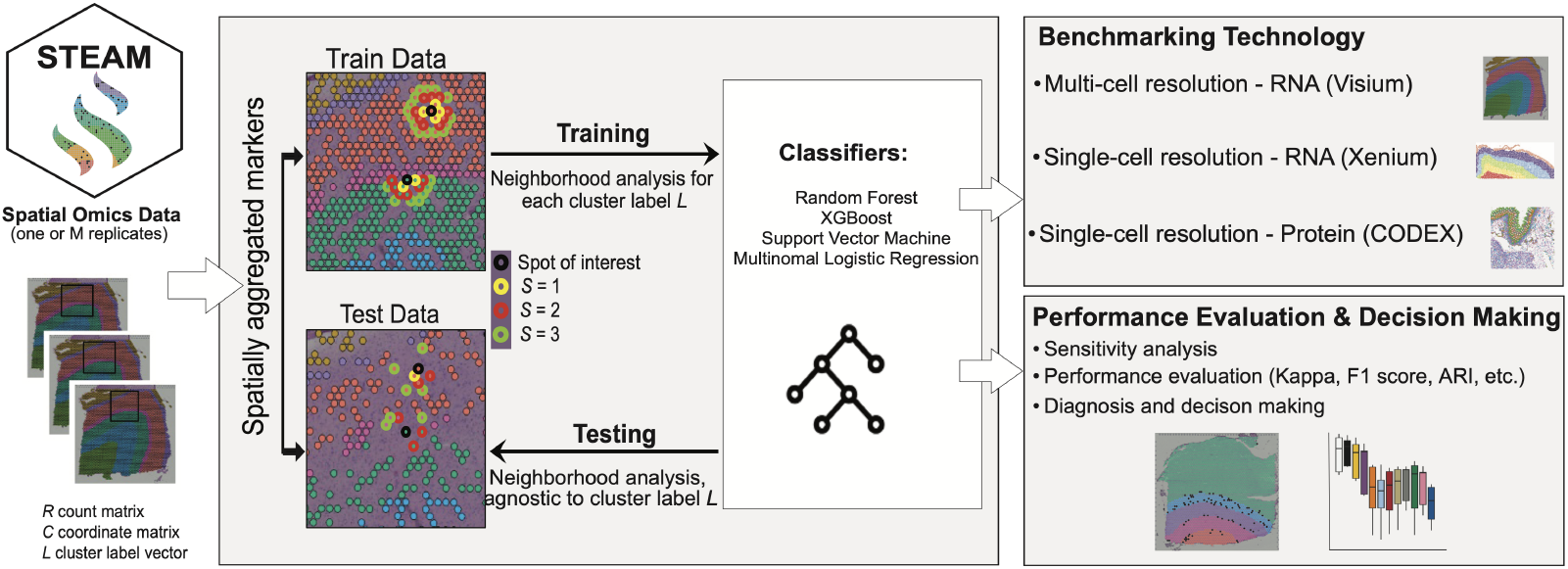
An overview of the STEAM framework and its systematic approach. STEAM processes spatial transcriptomic data (single- or multi-sample) with count matrix *R*, spatial coordinates *C*, and pre-identified cluster label *L*, assessing reliability via machine learning-based classification and prediction consistency. STEAM was benchmarked across different spatial platforms using various statistical metrics.

STEAM can be applied to various types of spatial omics, including multi-cell and single-cell resolution spatial transcriptomics, as well as spatial proteomics. Accordingly, STEAM takes in the same four inputs:

1. A normalized gene expression matrix *R* of size *n × m*, where *n* is the number of genes or proteins and *m* is the number of spots or cells present in a dataset.
2. A matrix *C* of size m^3^, where m represents the number of spots or cells, with 3 columns for the cell/spot names, *x*-coordinates, and *y*-coordinates.
3. A vector *L* of size *m*, where each entry corresponds to the cluster label or annotation for the cell or spot at that index.
4. A single integer *S* that specifies the neighborhood size to be averaged, a process detailed in the “Training data preparation” section.

### Spatially aggregated marker identification

We identify spatially aggregated markers to assess global spatial autocorrelation in gene expression patterns, as genes often exhibit localized expression that corresponds to distinct functional regions within tissues. Specifically, we use Moran’s I (Miller et al., 2021) to detect these spatially organized genes:

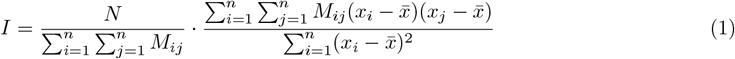

where *x*_*i*_ represents the expression value of gene *i* at a specific cell or spot in *R*, with *n* representing the total number of cells or spots. *M* is the spatial adjacency matrix constructed using coordinates in *C* and the neighborhood size *S*, with *M*_*ij*_= 1 if cells *i* and *j* are within distance *S* and *M*_*ij*_= 0 otherwise.

STEAM further refines Moran’s I by excluding patterns influenced by small cell counts, emphasizing the detection of localized clusters of spatially heterogeneous gene expression within each cell neighborhood using the Local Indicator of Spatial Association (LISA) (Anselin, 2010). For each cell *i*:

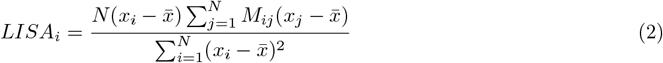

This process yields a new matrix where p represents the number of genes that meet the spatial heterogeneity criteria. Genes are filtered for robust spatial patterns, retaining only those for which at least 5% of cells exhibit a significant LISA score (p-value *<* 0.05).

### Training data preparation and neighborhood analysis

Following the formatting and inputs of the data, STEAM performs further processing steps prior to machine learning classification. First, the data is split into training and testing sets, with 70% of spots or cells assigned to training and 30% to testing. Following this separation, the training data is further divided based on cell type label. Each cell or spot in each label category then has its count data averaged across its spatial neighbors according to the inputted value of S. For example, if *S* is set to 3, the value of spot *i* is averaged across all spots with the same cell type label within 3 neighbors distance on the *x−* and *y−* coordinates informed by *C*. A visual of this process is available in Figure 1.

Specifically, we define:

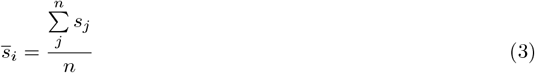

where *n* is the number of cells or spots with the same cell type label within the *S*-neighborhood coordinates of *s*_*i*_.

### Model building

After averaging each spot or cell within its neighborhood, the resulting expression values and corresponding labels from *L* are used to train a classifier. To achieve robust classification performance, we applied four different classification models, including tree-based methods like Random Forest and Extreme Gradient Boosting (XGBoost), SVM with various kernel functions, and a multinomial model.

We utilized the R package “caret” (version 6.0-94) to streamline the model building process. as it contains tools for data splitting, model selection as well as variable importance estimation. The Random Forest classifier was tested using various tree counts, finding that 100 trees achieved the most reliable accuracy while remaining computationally efficient.

For SVM, we conducted it with different kernel functions, including linear, radial basis function (RBF), polynomial, and sigmoid; employing the default parameters for each. Among these, the RBF and linear kernels yield the highest classification accuracy on the DLPFC, Xenium, and CODEX datasets.

For XGBoost, we implemented the “xgbTree” method in the “caret” package to capture complex, non-linear relationships in the data, leveraging its flexibility and computationally efficiency.

For the multinomial model, we utilized the “multinom” method We increased the maximum number of weights (MaxNWts) from the default value of 1,000 to 5,000 to accommodate the complexity of all datasets tested, which included the DLPFC, Xenium and CODEX datasets with multiple outcome labels. We retained the default maximum number of iterations (maxit) at 100, as it was sufficient for model convergence.

### Testing data preparation

As described in the “Training data preparation and neighborhood analysis” section, 30% of the spots or cells in the input count data are held out for testing. Unlike in the training data, where spot or cell counts are averaged by neighbors of the same cell type label, the testing data is averaged agnostic to label. This prevents data bleed that could inform testing data classification. Thus, each spot or cell in the testing data has its count data averaged based only on neighbors within the *S* spatial coordinate distance, regardless of the label identity of those neighbors. This process, along with its distinction from the training data, is illustrated in Figure 1.

## Dataset

To evaluate STEAM’s performance and robustness, we accessed and analyzed three publicly available datasets, each with well annotations for cell types and structures. First, we reanalyzed the dorsolateral prefrontal cortex (DLPFC) dataset (Maynard et al., 2021), which includes twelve different DLPFC slides with gene expression quantification using 10x Visium. Each slide was annotated into up to seven categories, which are also publicly available. Thus, each slide can be leveraged and evaluated independently and the whole DLPFC collection can also be aggregated to assess STEAM’s performance. The second dataset utilized was generated from 10x Xenium data of the mouse cortex, which is available at the 10x Genomics link (https://www.10xgenomics.com/datasets/fresh-frozen-mouse-brain-for-xenium-explorer-demo-1-standard). This dataset has been thoroughly analyzed using Seurat and reference data from the Allen Institute, providing reliable annotations (Yao et al., 2023, Hao et al., 2023). Lastly, we reanalyzed CODEX proteomics data of the intestine from the Human BioMolecular Atlas Program (HuBMAP)(Hickey, et al., 2023). This dataset was annotated with publicly available code from the MaxFuse paper by (Chen et al., 2024).

### DLPFC data processing

All twelve DLPFC slides were processed consistently using Seurat for spatial assays (Maynard et al., 2021, Hao et al., 2023). Specifically, each slide sample was loaded individually, and SCTransform was performed to 3,000 variable features to normalize and scale the data (Hao et al., 2023, Hafemeister, et al., 2019). Principal component analysis (PCA) was then performed to generate the first 50 principal components, followed by Uniform Manifold Approximation and Projection (UMAP) on the first 30 principal components to visualize expression diversity in 2-dimensional space (Hao et al., 2023, Becht et al., 2018). Next, the cell type annotations provided for each slide sample by Maynard, et al. were added to each Seurat object. Here, **Supplementary Table 1** lists the spot counts and identification numbers for each of the twelve slides.

To refine the 3,000 highly variable features identified for each slide into a more informed, robust gene list, we identified spatially variable genes using Moran’s I via Seurat (Hao et al., 2023, Paradis et al., 2019). Genes that were significant and had Moran’s I values over 0.15 in each slide were kept, resulting in an aggregated list of 185 genes. The expression matrices for each slide were then downsampled to include only these genes for downstream analysis and model evaluation.

### Xenium mouse cortex data processing

The analysis of the Xenium mouse cortex data was kept consistent with the methods outlined by Seurat (Hao et al., 2023). The “Tiny subset” was accessed from 10X Genomics and processed with Seurat (Hao et al., 2023) (10X Genomics). SCTransform was performed with 3,000 variable features, followed by PCA and UMAP on 30 principal components (Hao et al., 2023). The data was then cropped to isolate the cortical region, and cells were annotated using the Allen Brain Atlas as a reference for robust cell type decomposition (Yao et al., 2023, Hao et al., 2023, Cable et al., 2021). Subsequently, Seurat’s “BuildNicheAssay” function was used to define sections of the mouse cortex similar to those annotated in the DLPFC (Maynard et al., 2021, Hao et al., 2023). The resulting dataset includes 248 highly variable features over 10,570 cells. No additional gene subsetting was performed, as the number of features obtained was already relatively low.

### CODEX intestinal data processing

The CODEX data from HuBMAP was accessed and filtered to cells in the intestine and 48 relevant protein markers, following the approach outlined by the MaxFuse authors (Hickey, et al., 2023, Chen et al., 2024). Since the obtained data was already scaled, PCA was applied and then UMAP based on 50 principal components. Cell types were previously annotated in the provided data and were labeled accordingly. For clarity and power, various subtypes of enterocytes were combined into one single group based on biology interpretation. Although paired expression and chromatin data are available in the form of single-nucleus RNA-sequencing and single-nucleus assay for transposase-accessible chromatin with sequencing, we chose to evaluate how STEAM’s performance with protein-based spatial data alone. Furthermore, as the measured 48 protein markers were well established for the CODEX data, no further filtering was needed.

## Clustering Algorithms

### Seurat Clustering

Seurat-based clustering was conducted for each involved dataset using the “FindClusters” function in the Seurat pipeline (Hao et al., 2023). For each DLPFC 10x Visium slide, clusters were identified with a resolution of 0.8 on the normalized expression via regularized negative binomial regression model, following Seurat’s tutorial, using the first 30 principal components (PCs). For the 10x Xenium dataset, clusters were identified with a resolution of 0.3 according to the analysis outlined in the Seurat vignette guidelines (Hao et al., 2023), also using 30 PCs. Lastly, each of the four CODEX slides was clustered with a resolution of 0.8, with 30 PCs applied for clustering. All slides were subsequently analyzed with STEAM using the Seurat-based clusters as label inputs.

### BayesSpace Clustering

BayesSpace (Zhao et al., 2021) was developed using a Bayesian statistical model to enhance the resolution of spatial transcriptomic data for clustering. By incorporating a spatial prior, it encourages neighboring cell spots to cluster together, based on the assumption that nearby spots are likely to belong to the same group. We applied BayesSpace to each DLPFC sample by first converting the data into a “SingleCellExperiment” format, allowing for streamlined analysis with BayesSpace.The expected number of clusters was set based on the manual labels, in particular, 5 or 7 for the layers in the DLPFC dataset. We utilized the BayesSpace software (version 1.5.1) and adhered to the default settings recommended in the tutorial. As suggested, we used the first 15 principal components for clustering. The expected number of layers was set to 5 or 7 depending on the DLPFC sample to align with the known cortical layers that capture the brain layers and white matter region. Clustering was performed with 1,000 Markov-Chain Monte Carlo (MCMC) iterations, with an initial burn-in period of 100 iterations. After this, we applied STEAM to evaluate cluster consistency and robustness using the BayesSpace clusters as label input.

### DR.SC Clustering

DR.SC (Liu et al., 2022), dimension-reduction spatial-clustering, was a method built through simultaneous dimension reduction and spatial clustering. A latent hidden Markov random field model was employed to enforce the spatial smoothness in clustering. We used the DR.SC method to analyze the DLPFC slides. For the DLPFC slides, the expected number of clusters was provided according to the manual label classifications, as with BayesSpace (5 or 7 depending on the sample). Specifically, we used default parameters as detailed in the official tutorial on the DR.SC software’s (version 3.4) Github for DLPFC data analysis, which provides a comprehensive guide for data analysis Upon this, we applied STEAM with the DR.SC cluster labels as input.

### SpaceFlow Clustering

SpaceFlow (Ren et al., 2022) was a computational tool developed using spatially regularized deep graph networks to framework to integrate gene expression with spatial information to generate spatially aware embeddings. These embeddings serve as the basis for deriving domain segmentation and a pseudo-Spatiotemporal Map (pSM), effectively capturing spatiotemporal patterns and characterizing the spatial-aware clusters in tissue. We used SpaceFlow to analyze the DLPFC slides. Specifically, all model parameters were set to their default values as specified in their Github documentation. The Deep GraphInfomax Model was optimized with a set of hyperparameters for the training step. Spatial regularization strength was set to 0.1, the number of latent embeddings to 10, and the learning rate was set at 1e-3. Training regimen spanned up to 1,000 epochs, with a max patience of 50 and a minimum stop at 100, following recommended setting.

## Performance Metrics

F1 score: a measure of predictive performance and is considered the harmonic mean of precision and recall as it balances the trade-off between these two metrics and is useful when assessing imbalance datasets. F1 score is calculated as follows:

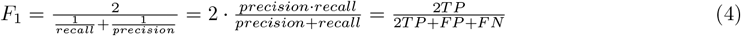

Where precision is the number of true positives divided by the number of all predicted positives and recall is the number of true positives divided by the total number of true positives and false negatives.

Kappa score (or Cohen’s kappa): a statistical measure to assess the performance of multi-class classification models. Unlike simple accuracy, the kappa score adjusts for imbalances in class distribution, making it a more robust metric. For each model, Kappa is the primary statistic that is output in the console for model evaluation and is used alongside other metrics to comprehensively assess model performance. Cohen’s kappa is calculated as:

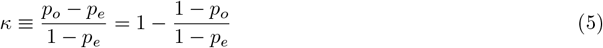

where *p*_*o*_ is the observed agreement that represents the proportion of instances where the raters or classifiers are in agreement. *p*_*e*_, the hypothetical probability of agreement by chance, is based on the observed data and the probabilities of randomly selecting each category or class label.

ARI (Adjusted Rand Index): a statistical measure used to quantify the similarity between two partitions of a dataset. It evaluates how well the assignments of data points to clusters agree while accounting for chance by comparing the observed agreement to the expected similarity under a random model. The ARI value ranges from -1 to 1, where negative values indicate less agreement than expected by chance, 0 represents random agreement, and 1 indicates a perfect match between the partitions.

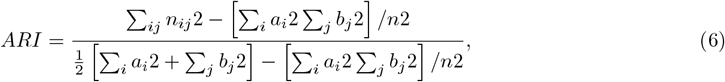

## Results

### STEAM achieves high accuracy for prediction of true cluster annotations on spatial transcriptomics

To evaluate the performance of STEAM, we implemented and benchmarked different machine learning models under STEAM framework using the DLPFC brain samples generated from 10x Visium data where manual annotations are available (**Figure 2A, see Supplementary material Dataset**). To identify the best-performing models and configurations for accurate cluster annotation, we benchmarked RandomForest (RF), SVM, XGBoost (XGB), and Multinomial Log Linear Models (MN), with manual layer labels provided by the original study as ground truth. For sensitivity analysis, we investigated the impact of varying neighborhood sizes on model performance by testing sizes of 1, 3, 5, 10, and 25 (**Figure 2B**). After quantifying the Kappa coefficient for each model, we found that STEAM achieved the highest Kappa with a neighborhood size of 5, 78.1% for STEAM RF, 80% for STEAM XGB, 82.3% for STEAM SVM, and 79.9% for STEAM MN across all samples (**Figure 2C**). Similarly, the neighborhood size of 5 achieved the best ARI, accuracy, and F1-score (**Supplementary Figure 1A-C**). Given this observation, a neighborhood size of 5 was used for further analysis to determine the performance of the four models. Notably, our analysis revealed that the STEAM RF, STEAM XGB, and STEAM SVM models provided more consistent results than STEAM MN, achieving higher F1 scores (0.73, 0.78, 0.78 vs. 0.48), improved kappa coefficients (0.78, 0.79, 0.8 vs. 0.6), and more stable ARI values (0.70, 0.71, 0.71 vs 0.59) (**Figure 2D-F, Supplementary Figure 1D**). Note that four of the twelve slides (IDs 151669, 151670, 151671, and 151672) exhibit relatively higher ARI (**Figure 2F**). This is due to their five spatially informed clusters, as opposed to the seven existing clusters in for the other slides. Further, we assessed the computational efficiency and found that most of the models are scalable except XGBoost (**Figure 2G**). Taken together, we demonstrated that RF and SVM are accurate, robust, and efficient, making them suitable for most of the analyses. Moreover, consistent with current spatial transcriptomic classification methods, STEAM may encounter the common challenge of accurately classifying spots along boundaries between classes, particularly in narrow layers like Layer 4, where cellular composition and expression profiles gradually shift (**Figure 2H**). Despite this challenge in the transitional spatial regions, STEAM accurately captures the overall structure of the DLPFC samples, indicating promising feasibility for application to data generated from the same platform.

**Figure 2.**
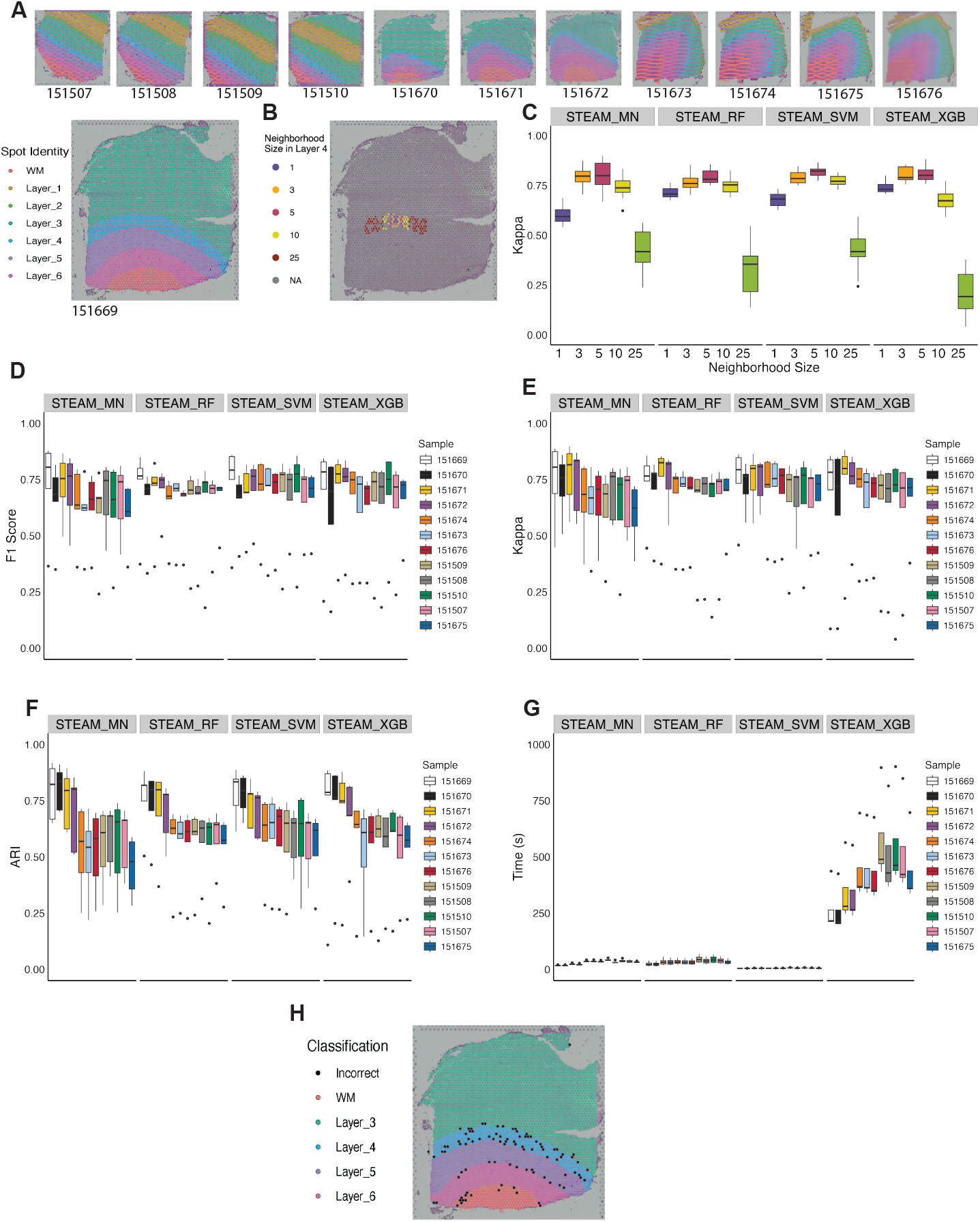
Comprehensive Analysis and Visualization of the STEAM Framework on DLPFC dataset.A. A visualization of the 12 DLPFC slides, with 151669 enlarged as an example for further visualization. **B**. A visualization of neighborhood averaging on the training data according to neighborhood sizes of 1, 3, 5, 10, and 25 on 151669 slide Layer 4. **C**. The Kappa coefficient of varying neighborhood sizes across four models: RandomForest (STEAM RF), radial Support Vector Machine (STEAM SVM), Extreme Gradient Boosting (STEAM XGB), and Multinom (STEAM MN). **D**. The F1 score, **E**. Kappa coefficient, and **F**. ARI for the performance on 1, 3, 5, 10, and 25 neighborhood averaging sizes across four different models: RandomForest (STEAM RF), radial Support Vector Machine (STEAM SVM), Extreme Gradient Boosting (STEAM XGB), and Multinom (STEAM MN). **G**. The runtime of STEAM on each DLPFC slide with each model. **H**. Visualizing misclassified cells on slide 151669 using STEAM RF.

### STEAM maintains high accuracy for multiple replicates

Although spatial transcriptomics data are frequently collected from multiple adjacent tissue sections or multiple subjects, most existing methods focus on analyzing a single sample at a time due to intra- and inter-subject variation. However, since sections from the same tissue condition could exhibit similar cellular compositions, we hypothesize that extending STEAM from single-sample clustering evaluation to a multisample training approach could enhance clustering accuracy and spatial domain classification. By leveraging shared cluster information across samples through STEAM, this approach could help overcome challenges posed by sample-to-sample variation. Taking multiple replicates as input, samples were grouped according to their shared qualitative characteristics in their feature and spatial profiles: Group 1 includes samples 151669-151672 (n=4), and Group 2 includes samples 151673-151676 (n=4). Within each group, common spatially aggregated genes were identified across all samples to standardize features for analysis. We applied a 10-fold cross-validation technique, combining 80% of cells from each sample within the group for training and 20% for testing (**Figure 3A**). In conjunction with the cross validation, we also ran with 5 different seeds to account for variability in data splitting and model initialization to ensure the robustness and reliability of classification performance. STEAM RF and STEAM XGB achieved similar performance across metrics, with comparable Kappa, F1-score, ARI values, and accuracy in all groups (**Figure 3B-3D, Supplementary Figure 2A**). Overall, the results of this multi-sample analysis suggest that STEAM provides robust and reliable predictions of ground truth clusters, which can be applied to evaluate multiple-sample spatial clustering results.

**Figure 3.**
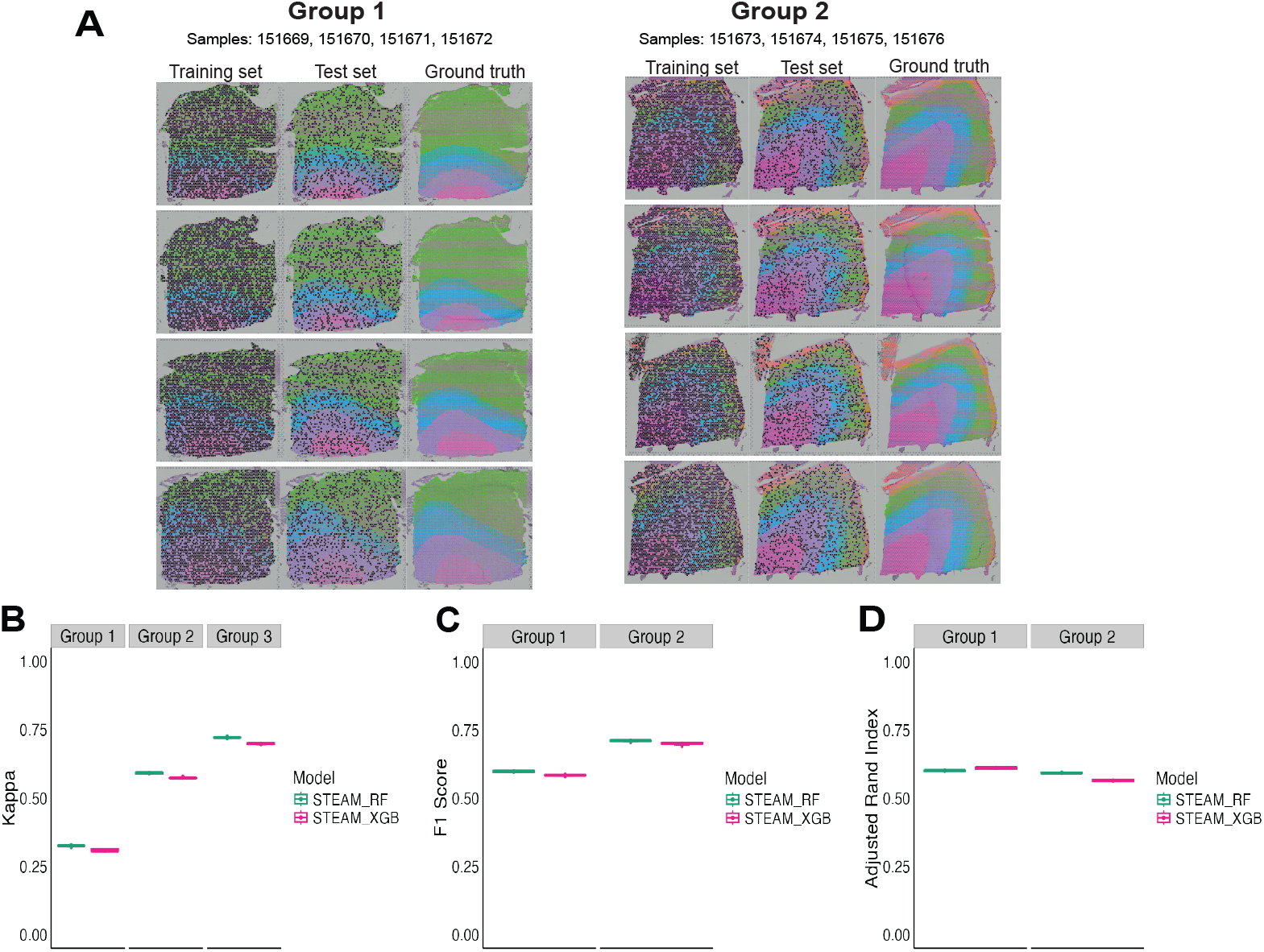
Multi-sample analysis and evaluation using STEAM. **A**. A visualization of the training, testing, and ground truth label for samples in each group for the multi-sample analysis. **B**. The Kappa coefficient, **C**. F1 score, and **D**. ARI of applying STEAM RF and STEAM XGB.

### STEAM is scalable to recover spatial niches on single-cell resolution Xenium data

Next, we evaluated the performance of STEAM using a 10X Genomics Xenium mouse cortex dataset with the aim of assessing how effectively the model handles single-cell resolution spatial transcriptomics, which provides finer spatial detail and more cells than 10x Visium. We utilized cell-type and niche annotations as ground truth, each offering a different level of specificity(**see Supplementary material Dataset**). Niche annotations group cells into six broader regions based on similarities in composition compared to the annotated 19 cell types (**Figure 4A**). Using 5-fold cross-validation with 5 random iterations across neighborhood sizes (1, 3, 5, 10, 25), we compared STEAM RF and STEAM XGB in terms of classification performance and computation time. STEAM RF consistently exhibited faster computation for both cell type annotation evaluation (**Figure 4B**) and niche annotation evaluation (**Figure 4C**). Despite its slower runtime, STEAM XGB slightly outperformed STEAM RF in classification accuracy. For both cell type annotation (**Figure 4D–F**) and niche detection (**Figure 4G–I**), STEAM XGB achieved slightly higher median scores across all metrics, including Kappa, F1-score, and ARI. We further assessed the impact of varying cell numbers on model performance, revealing that Kappa, F1-score, ARI, and accuracy remained stable across different randomly downsampled dataset proportions (40%, 60%, 80% and 100%) (**Figure 4J-O, Supplementary Figure 2D-E**). While STEAM RF and STEAM XGB achieved comparable classification performance in recovering cell types and spatial niches for high-resolution single-cell spatial transcriptomics, STEAM RF offered significantly greater computational efficiency, making STEAM RF more practical for time-sensitive analyses.

**Figure 4.**
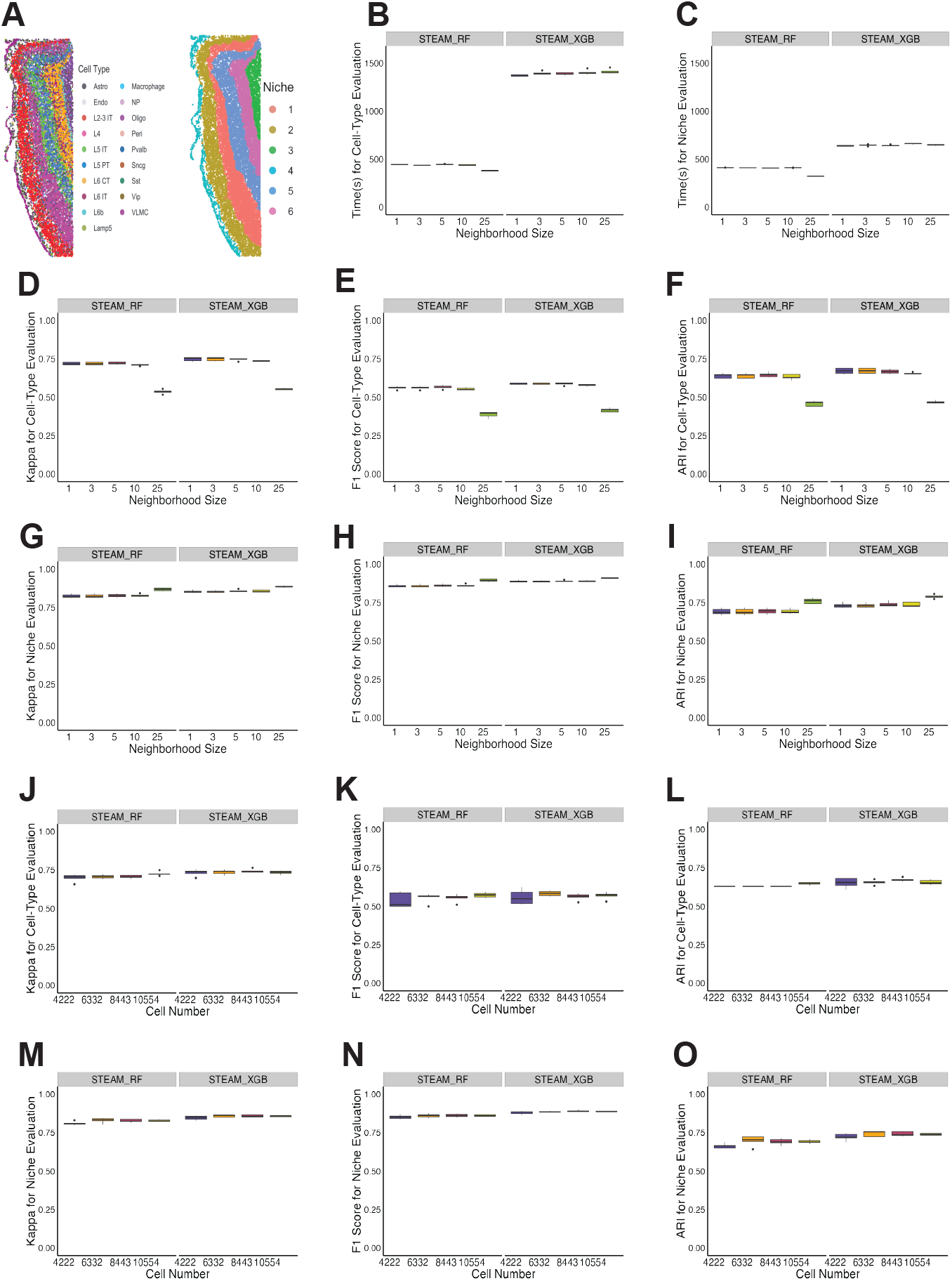
Benchmarking STEAM on 10x Genomics Xenium Mouse Cortex Data. **A**. A visualization of the Xenium dataset, with labels for both cell types and niches. **B**. The timing of STEAM RF and STEAM XGB for cell-type evaluation using different neighborhood sizes. **C**. The timing of STEAM RF and STEAM XGB for niche evaluation using different neighborhood sizes. **D**. The Kappa coefficient, **E**. F1 score, and **F**. ARI of evaluating Xenium cell type classification using STEAM RF, and STEAM XGB across 10 random iterations. **G**. The Kappa coefficient, **H**. The F1 score, and **I**. ARI of evaluating Xenium niche classification using STEAM RF, and STEAM XGB across 10 random iterations. **J**. The Kappa coefficient, **K**. F1 Score, and **L**. ARI of randomly sampled subsets of the data for cell-type evaluation. **M**. The Kappa coefficient, **N**. F1-score, and **O**. ARI of randomly sampled subsets of the data for niche evaluation.

### STEAM is robust to predict spatially informed cell types using spatial proteomics data on CODEX

Furthermore, we applied STEAM on a spatial proteomics dataset generated from the intestinal CODEX platform from HuBMAP (Hickey et al., 2023). Based on 48 well-characterized protein markers, this data revealed 15 cell types annotated by the original study displaying notable empty areas and unique architectures (**Figure 5A-D, see Supplementary material Dataset**). Unlike the more uniformly structured human brain, multiple cell types may be spatially co-localized with highly heterogeneous mesenchymal structures in the intestine, posing additional challenges for computational algorithms. We further visualized the instances of the misclassified cells by STEAM RF (**Figure 5E-H**), which, as expected, are primarily located at the boundaries between adjacent cell types. Benchmarking across all samples, STEAM RF and STEAM XGB demonstrated comparable classification performance, with a median Kappa of 0.87 vs. 0.89, a median F1-score of 0.64 vs. 0.67, a median ARI of 0.82 vs. 0.83, and a median accuracy of 0.89 vs. 0.91 (**Figure 5I-K, Supplementary Figure 2F**). Thus, this application and evaluation further highlight STEAM’s versatility in accurately predicting cell type labels from training data in heterogeneous spatial proteomics, which contains fewer molecular markers than that of spatial transcriptomics.

**Figure 5.**
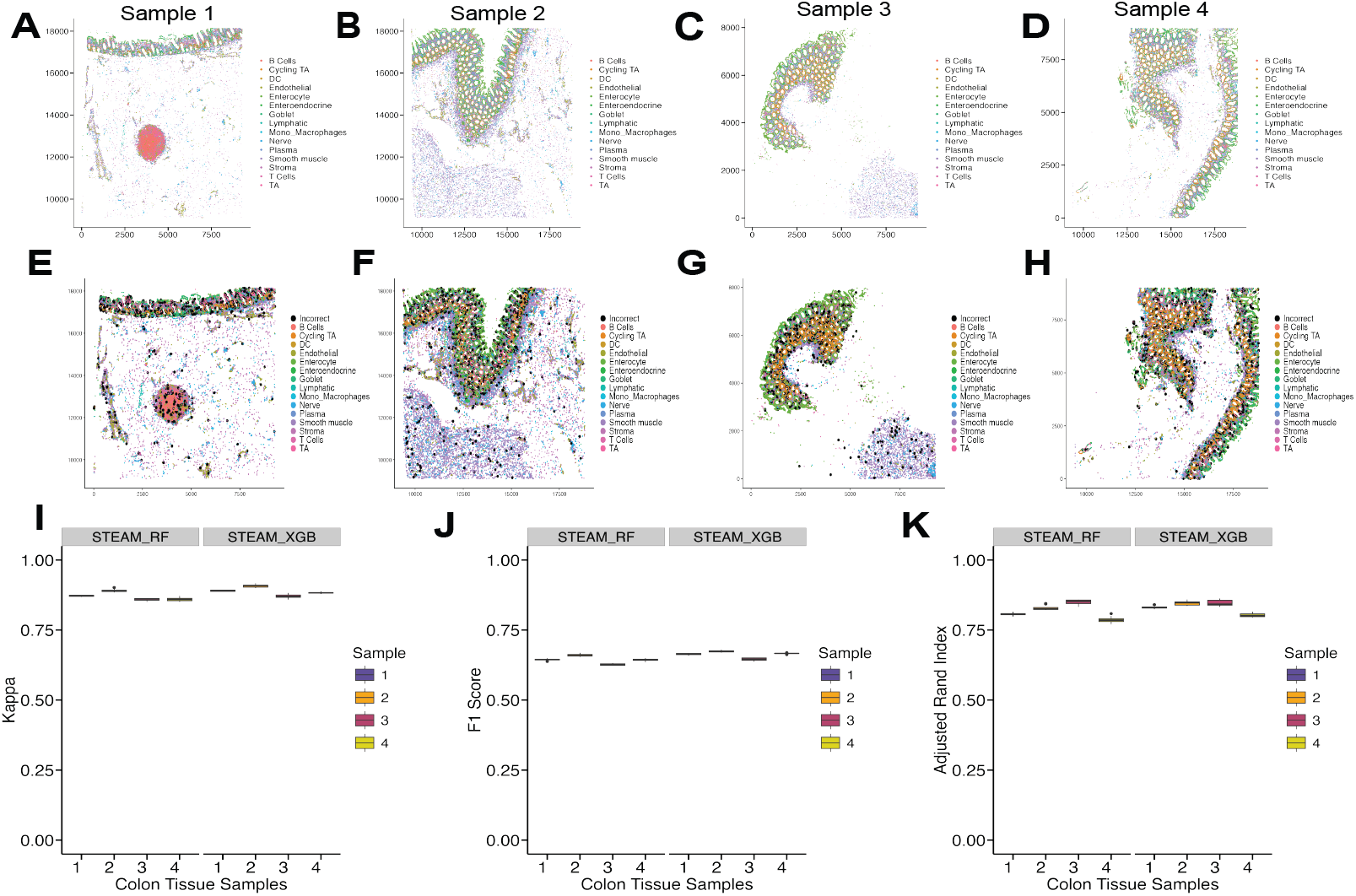
Benchmarking STEAM on CODEX intestinal data from HuBMAP. **A-D**. A visualization of cell types of the CODEX slide, with the annotation for all 4 slides on the right. **E-H**. A visualization of CODEX slide classifications, highlighting STEAM RF misclassifications. **I**. The Kappa coefficient, **J**. F1 Score, and **K**. ARI of STEAM RF and STEAM XGB on each CODEX slide across 10 random iterations.

### Demonstration of the usage of STEAM to evaluate clustering performance across multiple spatial transcriptomics analytical pipelines

Given the superior performance of STEAM in evaluating clustering accuracy and robustness using multiple benchmarking datasets (**Figure 2-5**), we aim to use STEAM as an effective framework to assess the performance of various clustering methods, providing investigators with detailed insights to guide decisions for downstream analysis. We selected clustering algorithms which represent different methodological categories: BayesSpace (Zhao et al., 2021) and DR.SC (Liu et al., 2022), built based on statistical modeling, and SpaceFlow (Ren et al., 2022) developed based on graph-based methodology, with all three considering both spatial and transcriptomics information. We also included Seurat which was chosen for its lack of consideration of spatial architecture in cluster assignment, serving as a benchmarking method. All four methods were tested across the 12 slides from the DLPFC (Maynard et al., 2021) dataset, using both STEAM RF and STEAM XGB with a 5-fold cross validation and 5 random seeds to account for variability in data splitting and model initialization for each sample. Consequently, BayesSpace and SpaceFlow outperformed DR.SC and Seurat in Kappa, F1-score, and ARI (**Figure 6A-C, Supplementary Figure 3A**). This was further supported by per-sample results, where the performance across multiple slides within each method demonstrated consistent outcomes (**Figure 6D-E, Supplementary Figure 3B-E**). Overall, the results from STEAM align with the computational model’s design, where spatially informed clustering tools, BayesSpace and SpaceFlow, outperformed other spatial-ignorant methods. Thus, STEAM offers a promising approach to guide researchers for decision making and model improvement for spatial omics clustering robustness.

**Figure 6.**
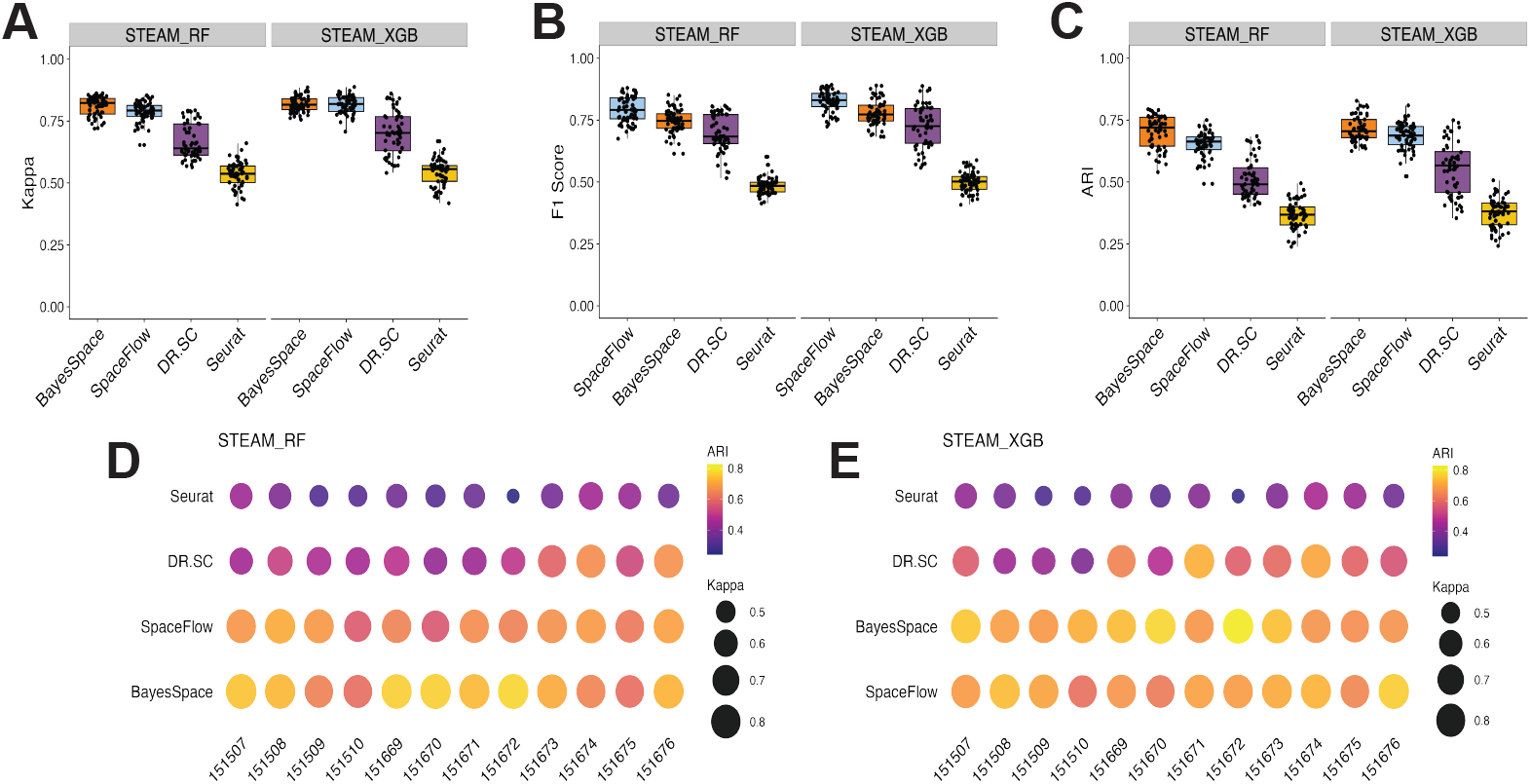
Benchmarking STEAM on various spatial transcriptomics clustering pipelines. **A**.Kappa coefficient, **B**. F1 Score, and **C**. ARI of the performance of using STEAM RF and STEAM XGB on the DLPFC slides, comparing clustering results from BayesSpace, DR.SC, Seurat, and SpaceFlow. **D**. The Kappa coefficients and ARI for each sample using STEAM RF. **E**. The Kappa coefficients and ARI for each sample using STEAM XGB.

## Discussion

Accurately evaluating clustering robustness and reliability of spatially coherent regions in gene expression and physical tissue structures remains a significant challenge in the absence of ground truth. This challenge is critical because reproducible clustering is essential for discoveries in spatial biology. To address this, we present STEAM, a comprehensive framework to assess the robustness of clusters and prediction accuracy for spatial omics data. By leveraging spatial proximity and covarying gene expression patterns, STEAM provides an objective and systematic approach to evaluate clustering methods and cell-type annotation consistency. We formulated our hypothesis on the premise that robust and consistent clustering, defined by coherent gene expression and spatial proximity, should enable accurate predictions of the unobserved cells within a labeled cluster when a subset of cells is used. Using a systematic approach, we benchmarked STEAM on a diverse range of spatial transcriptomics and proteomics datasets, encompassing both single-cell and multi-cell resolution. This benchmarking highlighted STEAM’s capability to deliver accurate and reproducible insights, while offering practical guidance for researchers in their downstream analyses. Markedly, STEAM facilitates multi-sample training, allowing researchers to assess cross-replicate clustering consistency. This feature enables cross-validate results across multiple samples, providing a crucial tool for ensuring reproducibility.

Despite its strengths, STEAM has limitations. One notable challenge lies in accurate classification of transitional regions, such as narrow or ambiguous tissue layers where cellular composition and gene expression profiles gradually shift. This has been a common challenge in computational imaging and spatial analysis workflows. For example, our analysis revealed a reduction in performance along boundaries in Layer 4 in the DLPFC dataset, likely due to the inherent heterogeneity and overlap of neighboring clusters. Future work could potentially focus on improving STEAM’s sensitivity to these transitional regions by integrating weighted models that explicitly account for these spatial gradients at cluster boundaries. Additionally, incorporating gene expression gradients into the framework could refine prediction accuracy, particularly in regions with high cellular and molecular heterogeneity. With the rapid increase in spatial omics data generation, STEAM is well-positioned to serve as an effective evaluation framework and one of the very first tools for benchmarking results. By enabling the assessment of clustering accuracy and reliability, we expect that STEAM could play a practical and promising role in driving robust and reproducible discoveries in spatial biology.

## Code Availability

The source code and the package are available at: https://github.com/fanzhanglab/STEAM.

## Funding

This work is supported by the PhRMA Foundation, Arthritis National Research Foundation, and the Translational Research Scholars Program award (F.Z.). We also acknowledge support from the NIH NLM Grant T15LM009451 (S.R.).

## Competing Interests

The authors declare no competing financial or nonfinancial interests.

## Author contributions statement

S.R. and C.S. implemented the method, analyzed the benchmarking experiments, and visualized the results. R.K. finalized the software package. S.R, C.S., and F.Z. wrote the initial draft. F.Z. supervised the study. All authors contributed to editing the final manuscript.

## Acknowledgments

We thank Madison Cho-Richmond for her insightful discussions on the manuscript. We also appreciate the Zhang Laboratory members for their valuable feedback and comments.

## Notes

### Competing Interest Statement

The authors have declared no competing interest.

