## Supplementary Materials for "STEAM: Spatial Transcriptomics Evaluation Algorithm and Metric for clustering performance"

| ID | Spot Count |
|---|---|
| 151507 | 4,221 |
| 151508 | 4,381 |
| 151509 | 4,788 |
| 151510 | 4,595 |
| 151669 | 3,636 |
| 151670 | 3,484 |
| 151671 | 4,093 |
| 151672 | 3,888 |
| 151673 | 3,611 |
| 151674 | 3,635 |
| 151675 | 3,566 |
| 151676 | 3,431 |

**Table S1.** The identification numbers and associated spot counts of each slide from the DLPFC dataset.

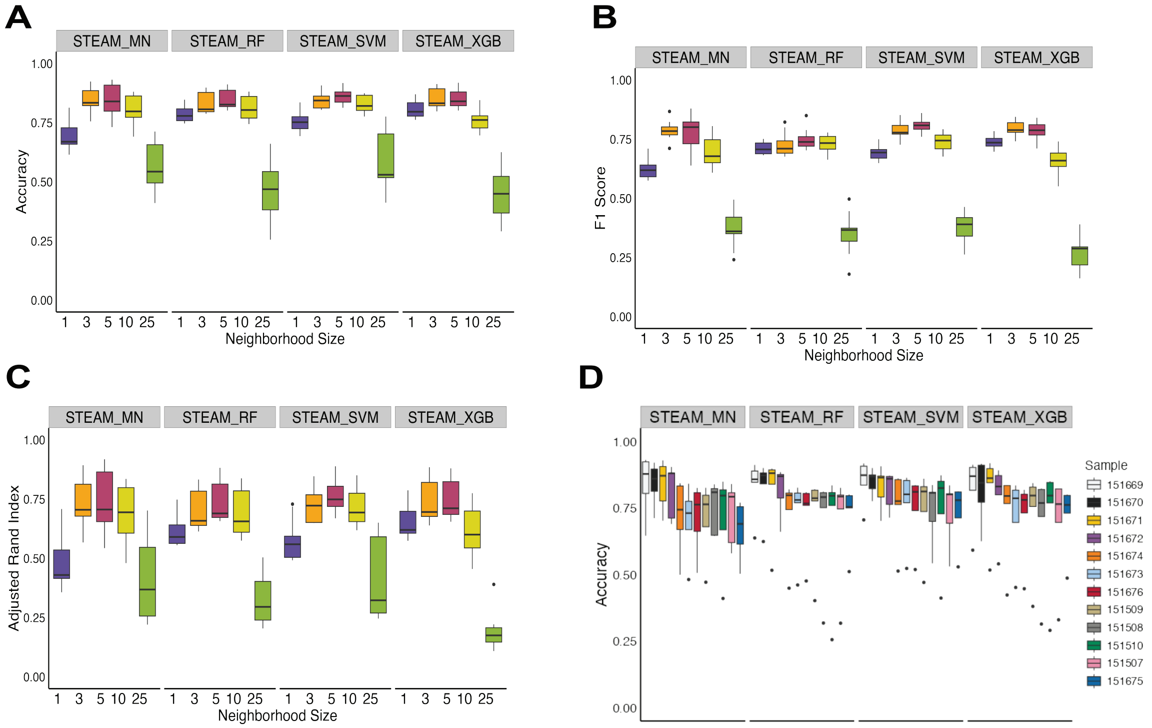


**Fig. S1. Evaluating model performance across neighborhood averaging sizes using DLPFC data. A.** The accuracy of 1, 3, 5, 10, and 25 neighborhood averaging sizes across four different models: RandomForest (STEAM_RF), radial Support Vector Machine (STEAM_SVM), Extreme Gradient Boosting (STEAM_XGB), and Multinom (STEAM_MN). **B.** The F1 score of 1, 3, 5, 10, and 25 neighborhood averaging sizes across four different models. **C.** The ARI of 1, 3, 5, 10, and 25 neighborhood averaging sizes across four different models. **D.** The accuracy of each DLPFC slide in each model across 10 random iterations.

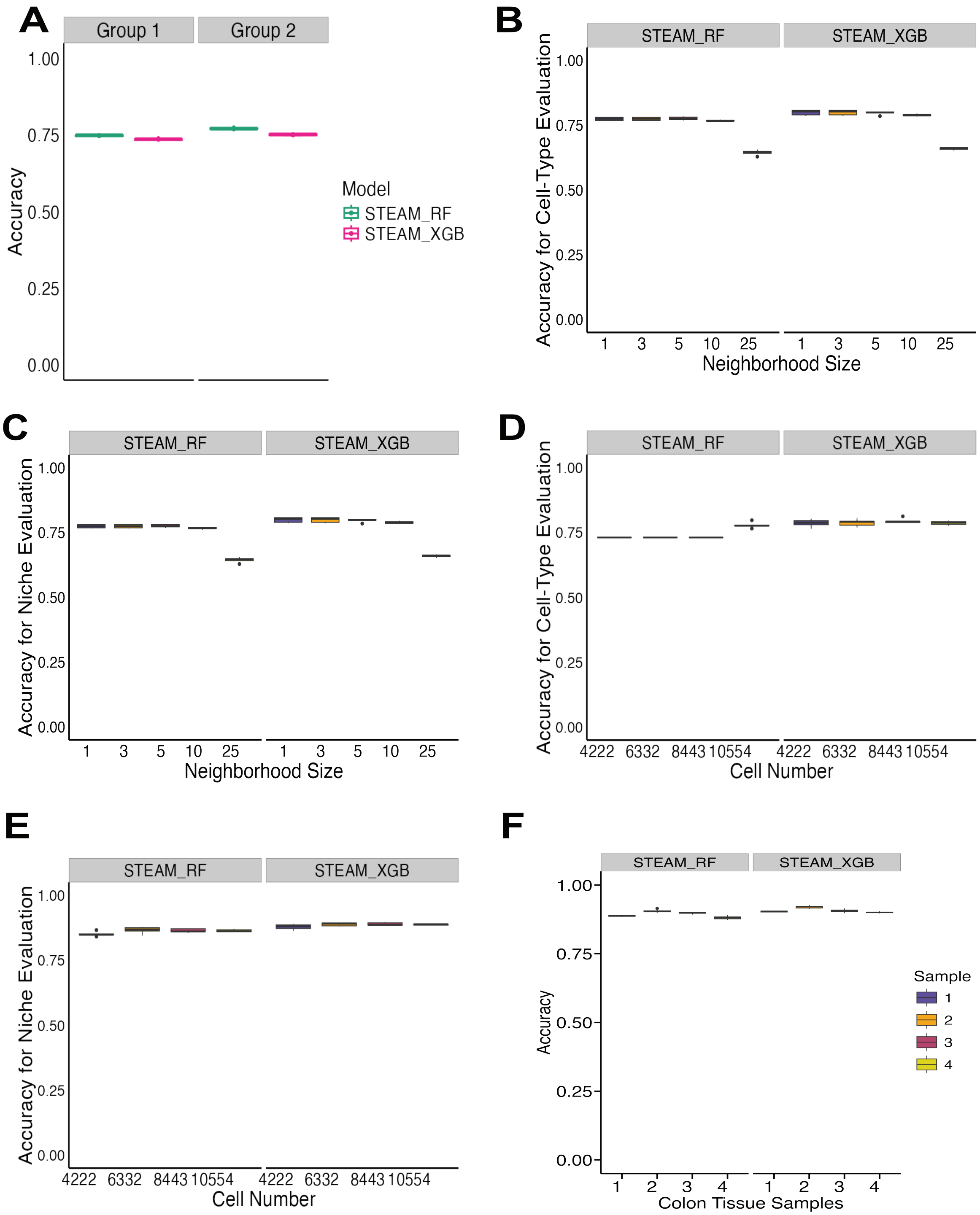


**Fig. S2. Additional metrics to evaluate model performance on the analysis of the DLPFC dataset, Xenium dataset, and colon dataset.** **A.** The accuracy of STEAM_RF and STEAM_XGB across the multi-sample groups. **B.** The accuracy of Xenium cell type classification using STEAM_RF and STEAM_XGB across 5 random iterations. **C.** The accuracy of Xenium niche classification using STEAM_RF and STEAM_XGB across 5 random iterations. **D.** The accuracy of using 40%, 60%, 80% and 100% randomly downsampled data from the Xenium dataset for cell-type classification. **E.** The accuracy of using 40%, 60%, 80% and 100% randomly downsampled data from the Xenium niche dataset for niche detection. **F.** The accuracy of STEAM_RF and STEAM_XGB on each of the 4 CODEX slides across 5 random iterations.

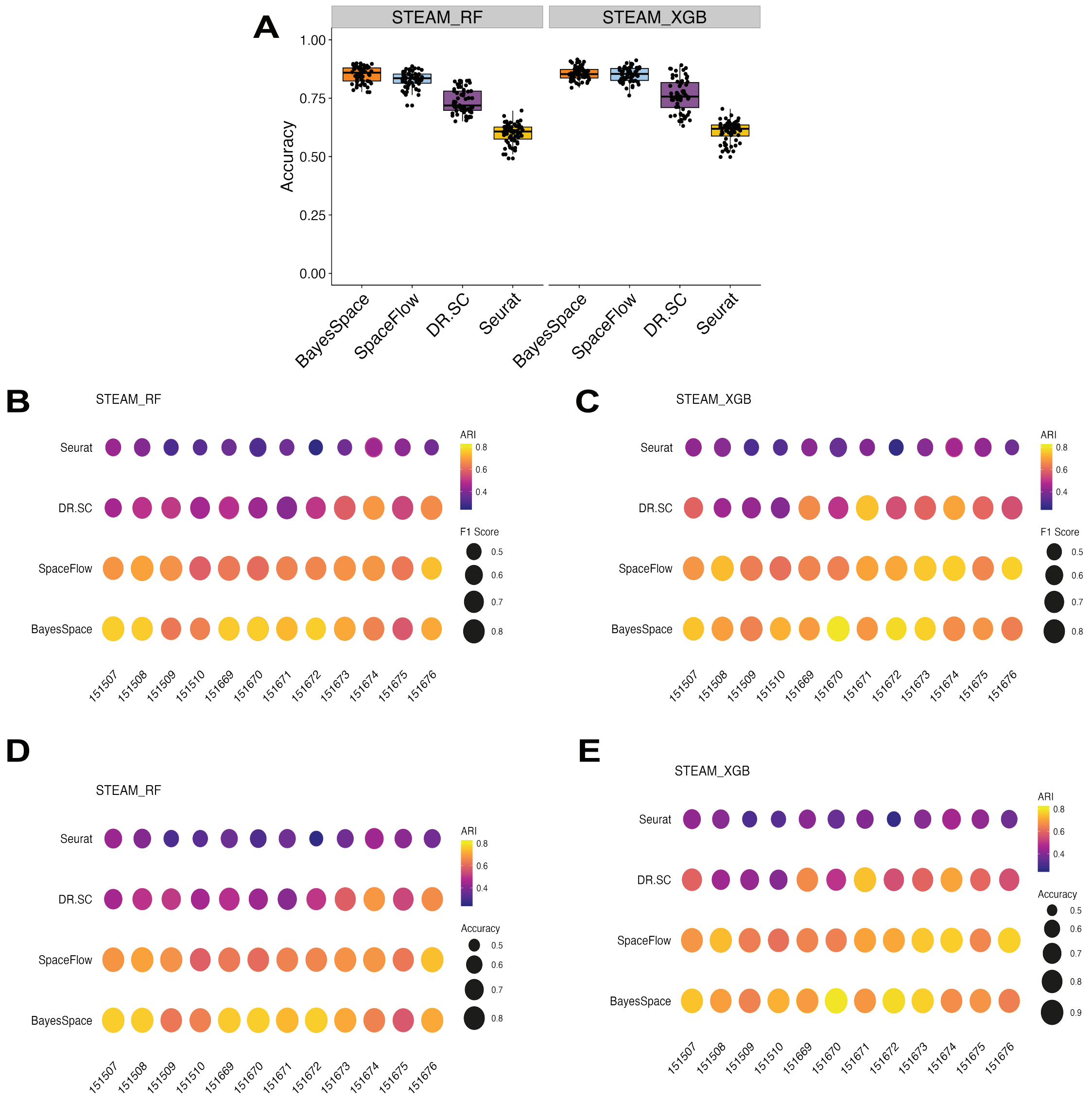


**Fig. S3. Comparative evaluation of various clustering algorithms. A.** Accuracies of STEAM_RF and STEAM_XGB on the DLPFC slides to evaluate clustering results from BayesSpace, DR.SC, Seurat, and SpaceFlow. **B.** The F1 score and ARI of STEAM_RF on the clusters generated from BayesSpace, DR.SC, Seurat, and SpaceFlow. **C.** The F1 score and ARI of STEAM_XGB on the clusters generated from BayesSpace, DR.SC, Seurat, and SpaceFlow. **D.** The accuracy and ARI of STEAM_RF on the clusters generated from BayesSpace, DR.SC, Seurat, and SpaceFlow. **E.** The accuracy and ARI of STEAM_XGB on the clusters generated from BayesSpace, DR.SC, Seurat, and SpaceFlow.
